# Seed preferences by ants and seed abundance determine associational effects between plant species in a semiarid grassland

**DOI:** 10.1101/2025.09.19.677408

**Authors:** García-Meza Diego, Martorell Carlos

## Abstract

Plant populations in semi-arid ecosystems rely heavily on seeds to endure environmental uncertainty. Granivores, like ants, cause significant seed loss but can also have a positive impact. Most studies on this topic examine seed consumption in isolation, failing to consider how the presence of other seeds can influence granivory. However, in nature, seeds of different species share space, which can lead to “associational susceptibility” or “resistance” effects. These effects depend on ant preferences for both the focal and neighboring species. Preferred seeds can act as a “magnet,” attracting ants and increasing the predation of nearby seeds. Conversely, less preferred seeds can have a “deterrent” effect. The magnitude of these associative effects may also depend on the preference of the focal seed itself: more desirable seeds will be less affected by neighboring seeds than less preferred ones. This study examined how the presence of heterospecific seeds affects ant seed removal in 28 plant species. We found that preferred seeds experienced a greater reduction in removal probability in the presence of other species. Additionally, plant population densities were positively correlated with the overall associative effect of neighboring seeds. This is the first study to report these indirect interactions between seeds affecting ant removal. Our results suggest that the impact of ants is a complex interaction of predation and dispersal, and their net effect on plant populations can be positive, even though seed removal is often considered a negative interaction.

## Introduction

Many plant populations from semiarid ecosystems rely heavily on seeds to endure environmental uncertainty (Brown & Ojeda, 1987). Granivores in these systems cause the loss of large amounts of seeds annually (Brown, Reichman, et al., 1979), but they can also exert a positive effect on plant populations through seed dispersal to suitable microsites (Arnan et al., 2010, 2012; Azcárate et al., 2005; Levey & Byrne, 1993; Retana et al., 2004). Thus, granivory (i.e., seed predation) in these systems could have large implications on population dynamics, community assembly, plant coexistence, and restoration and conservation plans (Brown, Davidson, et al., 1979; Brown, Reichman, et al., 1979; Davidson et al., 1985; Guo et al., 1995; Inouye et al., 1980; Sharafatmandrad & Khosravi Mashizi, 2021; Wills & Landis, 2018).

Despite the importance of granivory in desert ecosystems, there is a lack of information on how seed consumption by ants affects plant communities. Normally, granivory studies address how seed density, frequency, or seed characteristics affect seed removal considering each species in isolation. Nevertheless, seeds in nature share space with seeds of other species (Kiviniemi & Eriksson, 1999). Because the behavior of granivores, associational susceptibility or resistance could arise (Sotomayor & Lortie, 2015). Associational susceptibility occurs when removal rates are increased in the presence of companion species that attract a predator (i.e., magnet effect) (Sotomayor & Lortie, 2015). On the other hand, associational resistance occurs when predation of a focal species is reduced by another prey species (i.e., deterrent effect). While the mechanism causing an associational effect could be part of a reciprocally interaction, the term “associational effect” itself describes the net, often one-way, impact of the neighborhood on the focal species (Barbosa et al., 2009). In contrast, indirect interactions differ with associational effects regarding reciprocity (eg. (-/-), (+/+); Wootton, 1994). Indirect interactions caused by granivores have been studied mainly for rodents granivores in desert ecosystems (Veech, 2000, 2001), but we know little about the role of granivorous ants in mediating such interactions.

Associational effects may depend on ant preferences both for the focal species (the one whose predation is being analyzed) or the companion species (the one that affects the predation of the focal species). Companion species preferred by ants may be magnets by promoting trail formations toward the seed patch, increasing the removal rates of other species in their vicinity. Companion species with low preference could have a deterrent effect, acting as distractors (García-Meza et al. 2021). The focal species’ seed preferences could dictate the magnitude of the associational effects experienced. More preferred seeds will be less affected by companion species than less preferred seeds. This could be because, when favorite resources are being foraged, ant’s may be less distracted by other resources in the vicinity. Because ants are density-dependent foragers, it’s also possible that a plant’s abundance dictates preferences and thus how strong these associational effects are.

Associational effects may significantly impact plant populations. A deterrent effect can boost population densities by reducing seed predation, while a magnet effect can conversely decrease them by increasing losses to granivory. Critically, we often assume seed removal leads to predation, but ant-mediated seed dispersal can be beneficial, enhancing plant populations. For instance, ants may improve the population sizes of some species seemingly by dispersing them to appropriate sites, indicating that ant-plant interactions can be more advantageous than typically assumed (García-Meza & Martorell, 2022).

In this study, we aim to elucidate in 28 plant species whether seed removal by ants is affected by the presence of heterospecific seeds and if ant seed preferences mediate these effects. We expect that species that have high preferences will increase the seed removal of other seed plant species by prompting trail formation, thus eliciting associational susceptibility. On the other hand, we expect that low-preference species will decrease the probability of removal of other seed species, creating associational deterrence. Furthermore, we tested if ants’ preferences for the focal species used in this experiment modulates the associational effects experienced by companion seed species. We expect that the removal probability seeds with higher preferences will be less affected by companions compared to low-preference seed species. Furthermore, one of our goals is to see if these interactions and the focal species preferences have implications for plant populations. We expect that, due to an increase in the removal probability of species experiencing higher associational susceptibility, these will have lower population levels. Also, as ants are density dependent foragers, we expect that species with higher population levels are the ones preferred by ants. As far as our literature review goes, this appears to be the first study to determine if associational effects mediated by granivorous ants lead to indirect interactions between different seed species, their mechanisms, and the implications for plant populations.

## Methods

### Study Site

The study site is located at a semiarid grassland in Concepción Buenavista, Oaxaca, México (17° 55′ 43′′ N, 97° 25′ 14′′ W, 2202 m.a.s.l.). It has a mean annual precipitation of 530.5 mm. Seed dispersal of most of the plants in the grassland starts in late November and ends in late December (García-Meza et al., 2021).

Two ant species are the main granivores in this grassland, *Pheidole tepicana* and *Pheidole sciophila*. These two species have very similar size and behavior (Wilson et al., 2003). Both species are omnivorous, but they are most frequently seen transporting seeds. They have clear preferences for some seed species (García-Meza & Martorell 2024). Furthermore, these ants disperse as much as 10 % of the removed seeds (García-Meza et al., 2024).

### Experimental design for ants’ preferences

We studied 26 different plant species (Table 1) differing greatly in their densities in the grassland and in the sizes of their seeds. Two species (*Heterosperma pinnatum* Cav. and *Zinnia peruviana* (L.) L.) had two different types of seeds, which were analyzed separately, making a total of 28 different kinds of seeds. These were offered to ants as they are dispersed in nature (e.g., whole caryopses in Poaceae or achenes in Asteraceae) or with other organs (bracts in many Poaceae). We conducted a cafeteria experiment, which consists of offering different food items to predators and recording how fast and how many of the items are being removed. For simplicity, we will use the terms “species” to refer to the different kinds of propagules and “seed” to all propagules, irrespective of whether they were true seeds, fruits, or more complex structures.

**Table 1.** Species used in the cafeteria experiment.

| Species | Family |
| --- | --- |
| *Aristida adscensionis* L. | Poaceae |
| *Bidens pilosa* L. | Asteraceae |
| *Crusea simplex* (Willd.) J.H. Kirkbr. & Wiersema | Rubiaceae |
| *Digitaria ternata* (A. Rich.) Stapf . | Poaceae |
| *Dodonaea viscosa* (L.) Jacq. | Sapindaceae |
| *Echeandia flavescens* (Schult. & Schult. f.) Cruden | Asparagaceae |
| *Euphorbia mendezii* Boiss. | Euphorbiaceae |
| *Fimbristylis dichotoma* (L.) Vahl | Cyperaceae |
| *Helianthemum glomeratum* (Lag.) Lag. | Cistaceae |
| *Heterosperma pinnatum* Cav. | Asteraceae |
| *Ipomoea capillacea* (Kunth) G. Don | Convolvulaceae |
| *Microchloa kunthii* Desv. | Poaceae |
| *Milla valliflora* J. Gut. & E. Solano | Asparagaceae |
| *Panicum hallii* Vasey | Poaceae |
| *Plantago nivea* Kunth | Plantaginaceae |
| *Richardia tricocca* (Torr. & A. Gray) | Rubiaceae |
| *Sanvitalia procumbens* Lam. | Asteraceae |
| *Schkuhria pinnata* (Lam.) Kuntze ex Thell. | Asteraceae |
| *Setaria parviflora* (Poir.) Kerguélen | Poaceae |
| *Sporobolus atrovirens* (Kunth) Kunth | Poaceae |
| *Tagetes lunulata* Ortega | Asteraceae |
| *Tagetes micrantha* Cav. | Asteraceae |
| *Thymophylla aurantiaca* (Brandegée) Rydb. | Asteraceae |
| *Tripogandra purpurascens* (S. Schauer) Handlos | Commelinaceae |
| *Tripogonella spicata* (Nees) P.M. Peterson & Romasch. | Poaceae |
| *Zinnia peruviana* (L.) L. | Asteraceae |

We placed 76 experimental stations in the field, each consisting of a small depression of 5 × 5 cm and 1 cm depth to avoid removal by wind. On top of the depressions, we placed a structure that consisted of a 5 cm tall chicken mesh as walls and plastic plates as roofs to prevent rain or other granivores, such as birds or rodents, from removing our seeds. We placed on the holes a piece of sandpaper to create some friction to wind, make seed count easier, and mimic the soil. We have used these same devices previously and have found with video recordings that all removal can be attributed to ants (García-Meza et al., 2021).

Two different treatments were applied to the experimental stations. First, we placed a low seed-density treatment in which we added five seeds of a single species. The second treatment consisted of a heterospecific treatment. Because it would have been extremely difficult to count all seeds of all species rapidly enough if all species were placed together in a station, each one had five seeds of 8 different species. Seed mixtures were defined by randomly dividing the 28 species into seven groups of four species each. We then formed 20 groups of eight species by mixing the seeds of every possible pair of four-species groups and placed one of these combinations of eight species in each station in the field. This ensures that every possible pair of species appears together in at least one of these combinations. These combinations of seeds were placed at 7 am at the experimental stations and left them there for 24 h, counting the remaining seeds every 2 h until sunset and one last time at 6 am. Every day the treatments were assigned randomly to each station, and all species were randomly assigned to the seven groups of four species so that the combinations were different. The daily subdivisions into seven groups were chosen from a thousand randomized subdivisions so that the number of times each species pair occurred in a station throughout the experiment was as even as possible across all replications.

### Statistical analyses

Ants’ preferences for seeds of the studied plant species were calculated with the Rodger’s index (Krebs, 1999). This index considers the number of food items removed from experimental stations and how fast these items are being removed. It is defined as the cumulative consumption curve *A*:

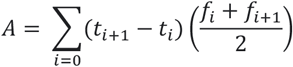

Where *t*_*i*_ is time elapsed since the start of the experiment, and *f*_*i*_ is the number of seeds removed in the *i*th observation. *A* becomes large when many seeds are removed rapidly, which indicates greater preference. We calculated the Rodger’s index using data from the low-density monospecific stations.

To estimate the effect of neighboring seeds on removal rates, we fitted the following model:

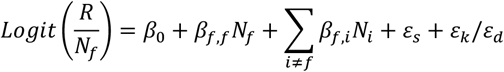

where *R* is the number of seeds removed of the focal species *f* at the end of each 2 h observation period, *N*_*i*_ is the number of seeds of species *i* at the beginning of each period, *β*_*f, i*_ is the effect of a seed of species *i* on the probability that a seed of the focal species is removed, and *ε*_*s*_ and *ε*_*k*_*/ε*_*d*_ . are the random effects of the experimental station and observation period nested in the day, respectively. The model was fitted with a binomial distribution in the package lme4 (Bates et al. 2015) in R (R Core Team, 2024). We adjusted the model for each species separately as focal, so we obtained a *β*_*f, i*_ value for each companion species affecting all 28 species. These *β*_*f, i*_ values for each companion species represent the change in seed removal of the focal species caused by seeds of each associated species. In other words, these values represent the sign and strength of the associational effect. Because this procedure was conducted to estimate the effect size of neighboring seeds rather than testing hypotheses, no model selection nor use of *P*-value was used to assess whether the effects of each companion species were significant. Instead, we recorded the standard error of *β*_*f, i*_ as an uncertainty measure of our estimate. Nevertheless, for 20 species, the full models had more support than the null one with a ΔAIC of two units, while 8 had the null as the best one (See Apendix 1 for detailed results).

With the estimated values obtained by these models, we arrange them to see if there are indirect interactions within each pair of species in the community. To do so, if the standard deviation of each *β*_*f, i*_ were greater than its absolute value, these effects were considered zero. This allows us to be cautious, as poorly estimated interactions will be accounted as no interaction.

To see if the associational effects between seeds of companion species on the removal probability depended on ants’ preferences of the focal and companion species, we performed a nested series of generalized additive mixed models (GAMM) with the package gamm4 (Wood & Scheipl, 2020) in R. The multiplicative inverse of the standard deviation of each *β*_*f,I*_ was used as weights in these analyses to account for the associational effects’ uncertainty. We then tested the focal and companion species identity as random effects on the model by adjusting them with restricted maximum likelihood and choosing the best one with at least two points in ΔAIC. We ended up using only the identity of the focal species as a random factor, and we then performed the analysis with maximum likelihood to evaluate for fixed effects. We selected the best model by two points in ΔAIC.

We then tested whether the indirect interactions between seeds and the preferences of ants influence the abundance of different species in the grassland. To do so, we regressed the mean abundance of the species in the study site over the past 20 years (See Zepeda & Martorell (2019) for details on how these data were obtained) on the mean effect of the interaction between companion species obtained in the past models and the Rodger’s index of each focal species with the package lme4 in R separately and compared them with their null model with a 2 -point difference in ΔAIC. The analysis with the mean effect of the interaction was made using the inverse sum of the variance as weights, while the analysis with the Rodger’s index was made using the inverse of the standard deviation of each species Rodger’s index. These measures helped the models fit, as they are a metric of uncertainty. Estimations that were more precise (i.e., had less uncertainty) had more weight in the analysis. As the relationship between Rodger’s index and the mean effect of the interaction with plant abundance could be sideways, we performed two weighted correlation analyses where we changed the response and explanatory variables of the past two described models. These were made to ensure that the weights adjusted in the explanatory variables in the linear models did not change the results when adjusted for the response variable. For both modeling procedures, this was the case (mean interaction model *P* = 0.051, mean interaction correlation *P* =0.051, preference model *P* = 0.53, preferences correlation *P*= 0.53).

## Results

Results of the associational effects of the 28 focal species used in our experiment can be found in Appendix 1. It was slightly more common that species have their removal probability reduced by their neighbors (15 out of 28 species, Figure 1 A), meaning that more than half of the species are experiencing associational resistance. Furthermore, most companion species reduce the removal probability of the focal species (18 out of 28; Figure 1 B), meaning that most species exert associational resistance within the community. For the 28 species analyzed, 20 had the full model (all companion species are considered) as the best one while 8 had the null one as the best model.

**Figure 1.**
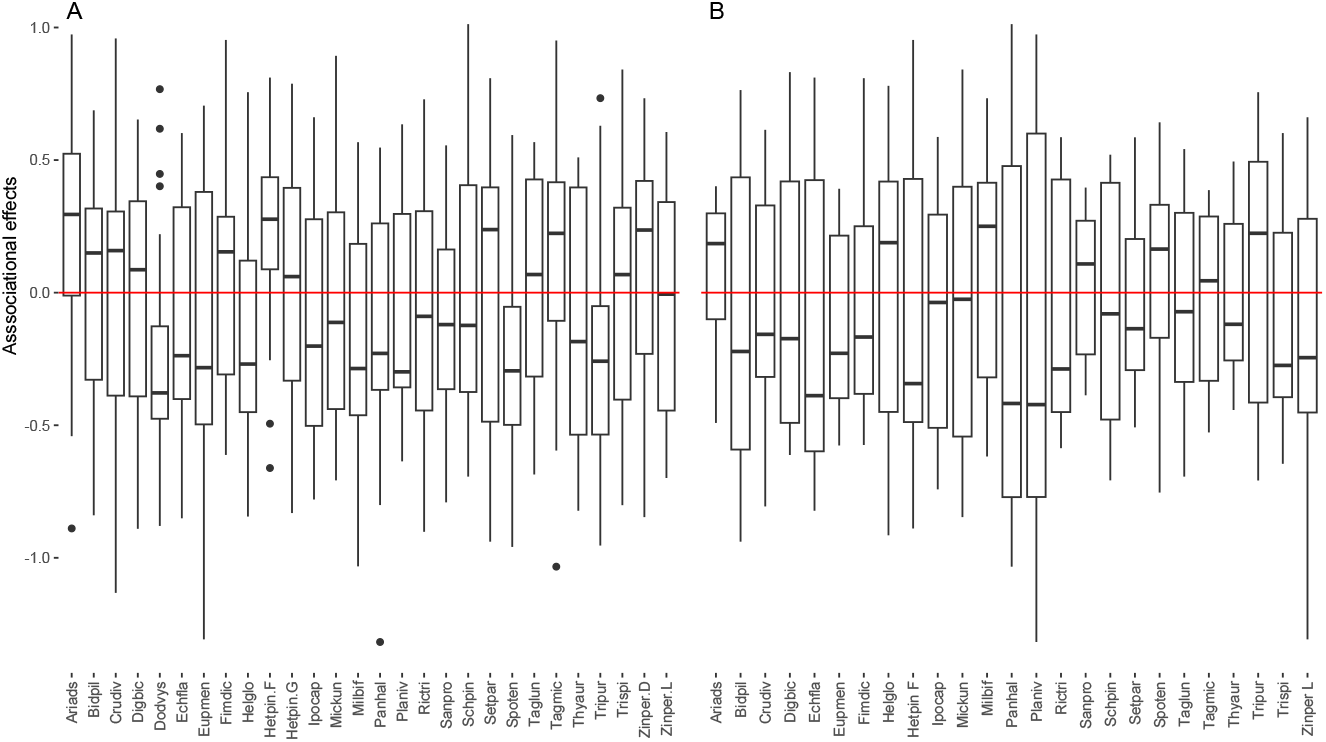
Mean change received (A) and exerted (B) on the removal probabilities for each species. Species are named after the first three letters of the genus and the first three letters of the species.

When interactions were arranged depending on their effect between species pairs, we found that the most common interaction in our community is a reciprocal augmentation on seed removal followed by a reciprocal reduction (Fig. 2). Apparently, non-reciprocal interactions between seeds are rare in the community.

**Figure 2.**
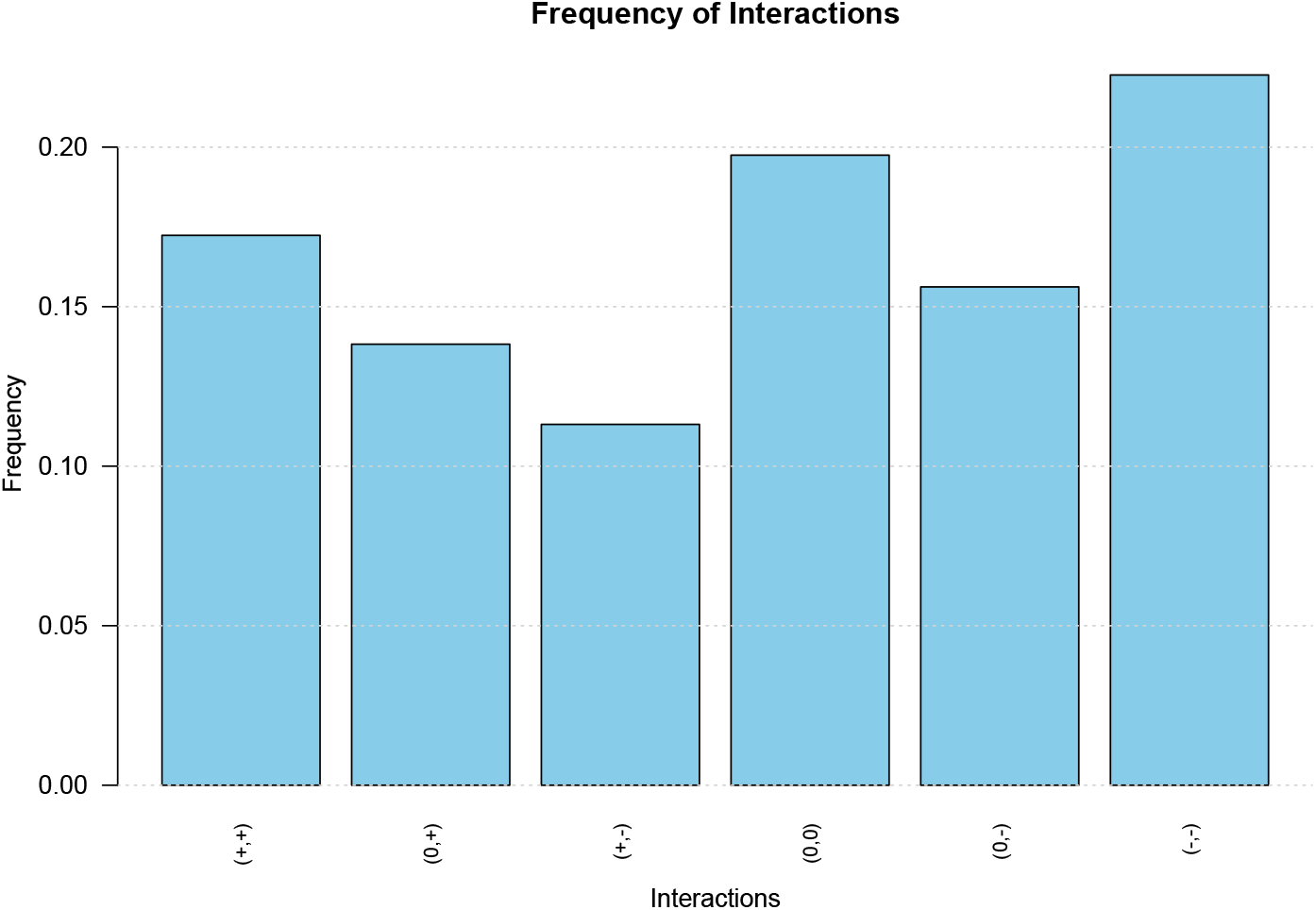
Frequency of reciprocal interactions found between species. Apparent competition (-,-) and indirect mutualism (+,+) are the inverse of the effects obtained in the model selection of each species as an increase in seed removal is linked with a reduction of the population levels.

Apparently, more preferred species suffered a reduction in their removal probability by the presence of other seeds in the neighborhood, while the least preferred species experienced an increase, regardless of how preferred the other seeds were (Figure 3).The generalized additive model selection results for the random effects showed that for all treatments and including and excluding interspecific interactions, the best model was the one that only included the focal species’ preference index, while the other models could be safely discarded ( ΔAIC > 2; Table 1).

**Figure 3.**
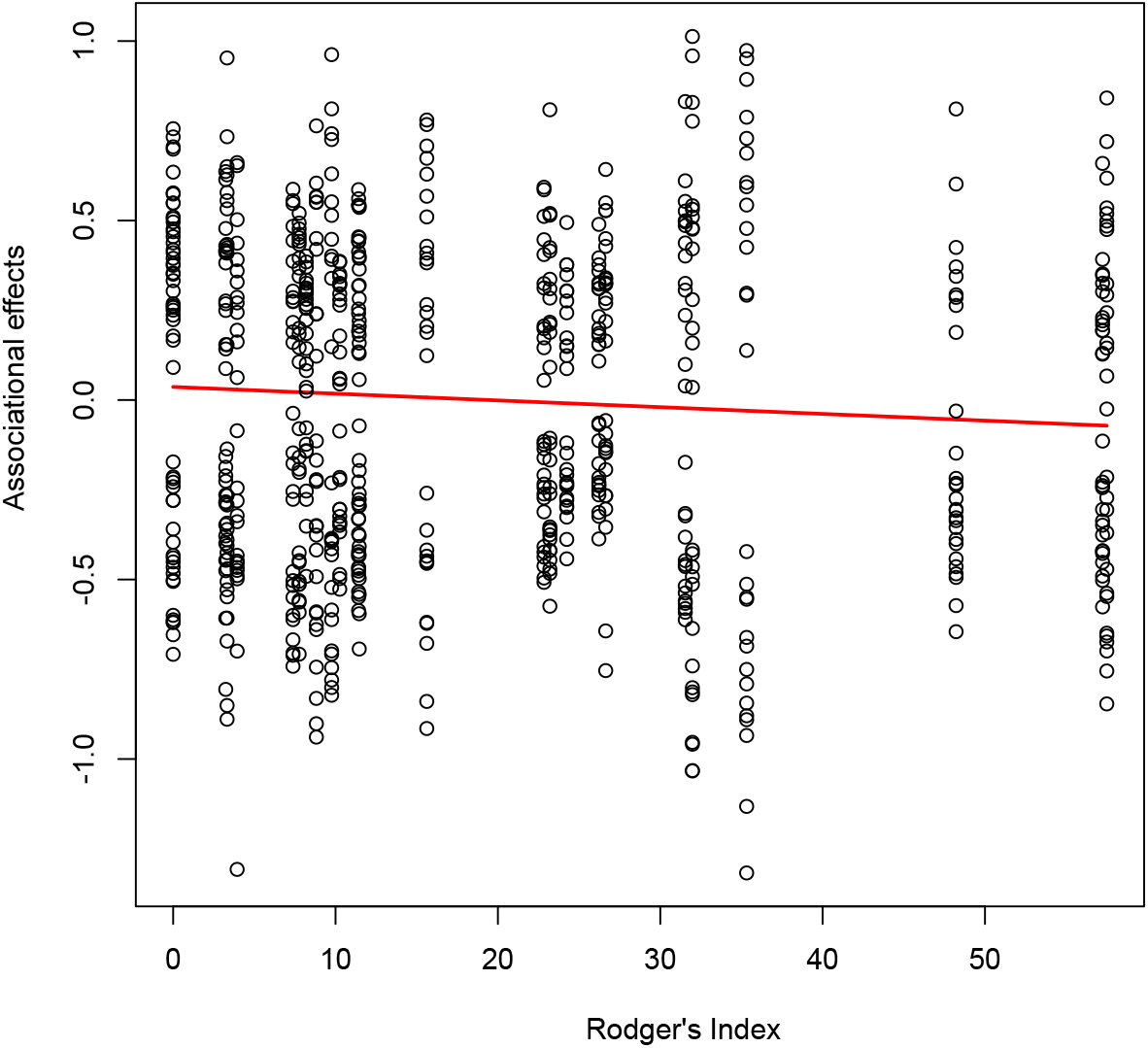
Relationship between focal species Rodger’s index (ants seed preferences) and the associational effects.

It would be expected that, without considering associational effects, preferred plant species (greater values on the Rodger’s index) would have the bigger population level as ants are density dependent foragers. Nevertheless, our analysis did not find this as the best model selected did not include the Rodger’s index as an independent variable (Δ AIC < 2). We also expected that population levels were related to the intensity of the associational effects a species is experiencing. Our analysis did find that there is a relationship between these two variables, and the best model to had a positive slope (Δ AIC > 2). This means that species experiencing an increase in their removal rate due to their neighbors, had bigger populations than those species with low or negative associational effects by the presence of companion species (Figure 4).

**Figure 4.**
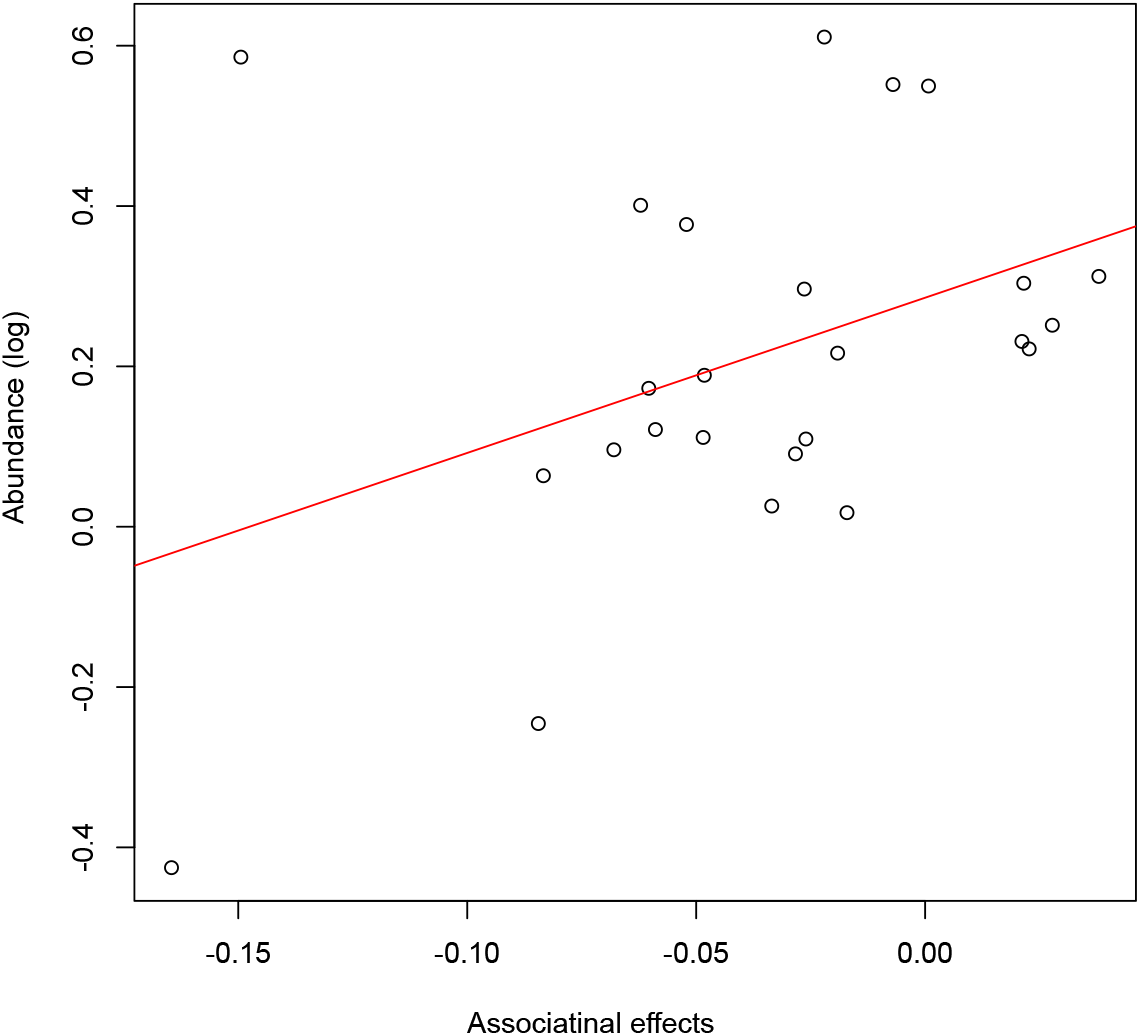
Relationship between associational effects on seed removal probabilities and the abundance of each species used in the experiment. The red line represents the model fitted.

## Discussion

In this study, we measured associational effects between different plant species’ seeds, mediated by ant seed removal. Results suggested that the presence of neighboring seed species influenced the removal rate of most focal seed species, either by increasing or decreasing it (Fig. 1 A and B). The most common interaction was a mutual increase in seed removal, followed by no interaction and then by a mutual reduction (Figure 4). Preferred seed species experienced greater reductions in removal probabilities by the presence of other plant species’ seeds (Figure 3). Apparently, these interactions had impacted plant populations in our community, as their densities were positively correlated with the overall associational effect of neighboring seeds on focal species’ removal probability (Figure 4).

The presence of other seeds significantly affected the removal rate of a focal seed species. Our findings show that associational resistance—a reduction in seed removal—is a more common outcome than associational susceptibility. This suggests that mixed-species patches might act as “safe sites” for seeds. This phenomenon could be explained by changes in ant foraging behavior. The presence of different seed types might increase ant handling times, reducing overall foraging efficiency. According to optimal foraging theory (Pyke et al., 1977), ants would then preferentially forage in low-diversity patches, which offer a more reliable reward for their effort.

The underlying mechanism for these indirect interactions may be linked to ant preferences. Our results indicate that seeds with low preference experience a “magnet effect” from neighboring seeds, while highly preferred seeds experience a “deter effect” that reduces their removal. This pattern could be a stabilizing mechanism for coexistence (Chesson, 2000). For example, it might prevent highly preferred species from being over-predated, though the population-level consequences of removal are still unclear. We found no correlation between ant preference and plant abundance. However, our analysis did show that more abundant species experienced a greater increase in their removal probability when in mixed patches. This suggests that rare species might avoid removal through these indirect interactions, while common species experience increased removal, which could act as a density-dependent regulation mechanism to prevent competitive exclusion (Chesson & Kuang, 2008).

An alternative perspective on this relationship between abundance and removal is that the causality is reversed: ants could be promoting the most abundant plant species. In this scenario, increased removal does not mean increased predation, but rather a greater probability of dispersal to favorable sites. This interpretation is consistent with a previous study in our community (García-Meza et al., 2024), which showed that ants increased the diversity of dominant species. Seed removal can lead to positive outcomes like dispersal to nutrient-rich ant nests and reduced intraspecific competition. Given the high density of ant nests in our study site, this dispersal-related help could be a powerful factor that drives the dominance of certain species. While the exact seed fate remains a topic of debate, our findings suggest that this complex interaction involves both predation and beneficial dispersal, which collectively influence plant community structure.

Beyond associational effects, we can also frame these dynamics as reciprocal indirect interactions between species pairs. Under this conceptualization, a pair of species that mutually reduce each other’s removal rates are considered indirect mutualism, while those that increase each other’s removal are engaged in apparent competition. Interestingly, when looking at these reciprocal interactions, apparent competition was the most common outcome, followed by no interaction and then mutualism. This contrasts with our findings for associational interactions, highlighting a key point: how we conceptualize and measure these indirect effects can dramatically change our understanding of their prevalence and impact on the community. Nevertheless, the average indirect effect across the community remains close to zero, suggesting that while specific interactions can be strong, the net community-wide effect is weak.

This study is the first to report indirect interactions between different seeds affecting ant removal. These interactions have significant implications for community dynamics and plant coexistence. The influence of ants on the community is not always negative, as is often assumed; instead, it appears to be a complex combination of predation and dispersal. While seed removal is typically considered a negative interaction, our results suggest that for some species, the net effect may be positive, mimicking the mutualistic interaction of myrmecochorous plants. Further research into seed fate and germination is needed to fully untangle this intricate plant-animal interaction.

