## Appendix 1 for "Seed preferences by ants and seed abundance determine associational effects between plant species in a semiarid grassland"

Results obtained with the generalized linear mixed models (glmm) for the effect of companion species on the probability of a seed to be removed of a focal species.

| Species | Model | df | AIC | ∆AIC |
|---|---|---|---|---|
| *Aristida adscensionis* | Full | 32 | 820.38 | 0.00 |
| | Null | 4 | 822.71 | 2.33 |
| *Bidens pilosa* | Full | 32 | 279.31 | 0.00 |
| | Null | 4 | 291.65 | 12.34 |
| *Crusea simplex* | Full | 32 | 446.59 | 0.00 |
| | Null | 4 | 458.45 | 11.87 |
| *Digitaria ternata* | Full | 32 | 708.92 | 0.00 |
| | Null | 4 | 792.52 | 83.59 |
| *Dodonaea viscosa* | Full | 32 | 192.99 | 0.00 |
| | Null | 4 | 203.51 | 10.53 |
| *Echeandia flavescens* | Full | 32 | 391.77 | 0.00 |
| | Null | 4 | 434.69 | 42.92 |
| *Euphorbia mendezii* | Full | 32 | 744.93 | 0.00 |
| | Null | 4 | 778.28 | 33.36 |
| *Fimbristylis dichotoma* | Full | 32 | 660.40 | 0.00 |
| | Null | 4 | 662.60 | 2.20 |
| *Helianthemum glomeratum* | Full | 32 | 568.30 | 0.00 |
| | Null | 4 | 652.54 | 84.24 |
| *Heterosperma pinnatum* | Full | 32 | 308.24 | 0.63 |
| | Null | 4 | 307.60 | 0.00 |
| *Heterosperma pinnatum* | Full | 32 | 289.85 | 0.00 |
| | Null | 4 | 303.93 | 14.08 |
| *Ipomoea capillacea* | Full | 32 | 277.52 | 9.95 |
| | Null | 4 | 267.57 | 0.00 |
| *Microchloa kunthii* | Full | 32 | 575.58 | 0.00 |
| | Null | 4 | 641.86 | 66.28 |
| *Milla valliflora* | Full | 32 | 366.28 | 0.00 |
| | Null | 4 | 414.87 | 48.59 |
| *Panicum hallii* | Full | 32 | 463.10 | 0.00 |
| | Null | 4 | 597.00 | 133.91 |
| *Plantago nivea* | Full | 32 | 501.62 | 0.00 |
| | Null | 4 | 670.99 | 169.37 |
| *Richardia tricocca* | Full | 32 | 525.48 | 0.00 |

| | | | | |
|---|---|---|---|---|
| | Null | 4 | 541.53 | 16.05 |
| *Sanvitalia procumbens* | Full | 32 | 804.82 | 4.08 |
| | Null | 4 | 800.74 | 0.00 |
| *Schkuhria pinnata* | Full | 32 | 480.68 | 9.10 |
| | Null | 4 | 471.58 | 0.00 |
| *Setaria parviflora* | Full | 32 | 661.57 | 0.00 |
| | Null | 4 | 673.57 | 12.00 |
| *Sporobolus atrovirens* | Full | 32 | 794.28 | 0.00 |
| | Null | 4 | 819.63 | 25.35 |
| *Tagetes lunulata* | Full | 32 | 595.60 | 6.22 |
| | Null | 4 | 589.38 | 0.00 |
| *Tagetes micrantha* | Full | 32 | 651.25 | 28.03 |
| | Null | 4 | 623.23 | 0.00 |
| *Thymophylla aurantiaca* | Full | 32 | 758.27 | 0.00 |
| | Null | 4 | 768.57 | 10.30 |
| *Tripogandra purpurascens* | Full | 32 | 354.52 | 3.60 |
| | Null | 4 | 350.92 | 0.00 |
| *Tripogonella spicata* | Full | 32 | 704.76 | 0.00 |
| | Null | 4 | 744.75 | 39.99 |
| *Zinnia peruviana* | Full | 32 | 121.91 | 0.00 |
| | Null | 4 | 141.32 | 19.41 |
| *Zinnia peruviana* | Full | 32 | 299.22 | 12.07 |
| | Null | 4 | 287.14 | 0.00 |
